# A Surgical Technique for Individual Control of the Muscles of the Rabbit Lower Hind-limb

**DOI:** 10.1101/2024.01.10.575091

**Authors:** Michael Baggaley, Andrew Sawatsky, Stephanie A. Ross, Walter Herzog

## Abstract

Little is known regarding the precise muscle, bone, and joint actions resulting from individual and simultaneous muscle activation(s) of the lower limb. An *in situ* experimental approach is described herein to control the muscles of the rabbit lower hind-limb, including the medial and lateral gastrocnemius, soleus, plantaris, and tibialis anterior. The muscles were stimulated using nerve-cuff electrodes placed around the innervating nerves of each muscle. Animals were fixed in a stereotactic frame with the ankle angle free to rotate in the sagittal plane to quantify the behaviour of the lower hind-limb muscles. To demonstrate the efficacy of the experimental technique, isometric plantarflexion torque was measured at a 90 ° ankle joint angle at a stimulation frequency of 100, 60, and 30 Hz. Individual muscle torque and the torque produced during simultaneous activation of all plantarflexor muscles are presented for four animals. These results demonstrate that the experimental approach was reliable, with insignificant variation in torque between repeated contractions. The experimental approach described herein provides the potential for measuring a diverse array of muscle properties which is important to improve our understanding of musculoskeletal biomechanics.

**Highlights:**

- A reliable surgical technique was developed for isolated activation of the plantarflexor muscles and the tibialis anterior in the rabbit. Joint torque data are presented for four rabbits at a single joint angle and three stimulation frequencies.

## 1. Introduction

Understanding how individual muscles function as part of the musculoskeletal system represents a fundamental problem in biomechanics (Crowninshield and Brand, 1981). The musculoskeletal system is composed of many components that interact with each other during muscle contraction. Little is known regarding the inter-muscular dynamics during contraction due to the difficulty in isolating individual components. Instead, much of our understanding of muscle function comes from experiments on isolated muscles; e.g., (Sleboda and Roberts, 2020; Siebert et al., 2016; Butterfield and Herzog, 2006). While critical to understanding basic muscle mechanics, these experiments are limited in their inferential capacity due to the removal from their operating environment. Instead, recent work has used ultrasound imaging; e.g., (Randhawa and Wakeling, 2018; Ryan et al., 2021) to retain *in vivo* muscle structure, thereby producing more accurate measures of inter-muscular dynamics. However, this approach only examines gross muscle function during voluntary activation, thereby limiting our ability to draw conclusions regarding the contribution of individual muscles to the collective behaviour of a muscle group.

Instead, a surgical procedure for individual muscle control was previously developed in our group to control the quadriceps muscles of the rabbit hind limb (Han et al., 2019, 2020; de Brito Fontana et al., 2018). Applying the surgical technique to the rabbit lower limb will expand our knowledge of the structure-function relationships that govern muscle. Myofascial connections differ between muscle groups, likely altering inter-muscular force transmission that can influence group behaviour (Finni et al., 2023). Functional differences due to muscle position, size, and orientation can be quantified, furthering our understanding of dynamic muscle function and the role of individual muscles within an agonist group.

This technical note describes a surgical and experimental approach to perform individual and simultaneous activation of agonist and antagonist muscles in the rabbit lower hind limb. This procedure allows us to quantify the role of each muscle in unique joint angle configurations, and how individual muscle action is altered by the simultaneous activation of all muscles. Data are provided to demonstrate the efficacy and reliability of the surgical technique.

## 2. Methods

### 2.1. Animal Preparation

Four skeletally mature New Zealand White rabbits were used to to develop the surgical approach and experimental protocol to test the behaviour of the medial and lateral gastrocnemius, soleus, plantaris, and tibialis anterior. All experimental procedures were approved by the Animal Ethics Review Committee of the University of Calgary (REB#AC20-0010).

### 2.2. Pre-Surgery Procedures

Prior to surgery, rabbits were injected with 1 mg/kg Acepromezine (Acevet 25 Injectable, Vetoquinol, Canada) and 0.02 - 0.05 mg/kg Buprenorphine (Vetergesic® Multidose 0.3 mg/ml Solution, Ceva Animal Health, UK). Rabbits were induced using 5 % isoflurane (Isoflurane USP, Fresenius Kabi Canada Ltd, Canada) to oxygen mixture while laying in a prone position. Throughout the experiment a surgical plane was maintained at a mixture of 2 -3 % isoflurane. Following anaesthesia, the animal was prepared for surgery by shaving the left hind limb and lower torso.

### 2.3. Surgical Procedures

Surgery was performed to implant a nerve cuff stimulating electrode on the nerves supplying the tibialis anterior, lateral gastrocnemius, medial gastrocnemius, and plantaris. The soleus was activated concomitantly with the laterial gastrocnemius, as the soleus nerve branch was not easily accessible. The soleus nerve bifurcates off the tibial nerve once it is deep to the lateral gastrocnemius; isolation of the soleus nerve would require significant damage to the lateral gatrocnemius muscle. Minimal alterations to muscle position and structure was sought to replicate the *in vivo* environment of the muscles.

#### 2.3.1. Tibialis Anterior Nerve Isolation

An incision (6-7 cm) was made on the posterior lateral aspect of the femur to access the peroneal and tibial nerve. Surface-level connective tissue and fat were removed using blunt dissection, revealing the underlying muscles. The biceps femoris and the semimembranosus were separated along their myofascial connection to gain access to the nerve. Once a cavity has formed between the biceps femoris and the semimembranosus, the peroneal nerve will be visible; it is the deepest nerve relative to the skin. The peroneal nerve was isolated from the surrounding connective tissue using blunt dissection, and a custom-built nerve cuff was attached. To ensure that only the tibialis anterior was activated with stimulation of the peroneal nerve, the remaining muscles that are also innervated by the peroneal nerve and cross the ankle joint were transected. To accomplish this, an incision (6-7 cm) was made on the lateral aspect of the tibia, just distal to the knee joint. Blunt dissection was used to separate the tibialis anterior and the lateral gastrocnemius along their myofascial connection, exposing the peroneal nerve. The peroneal nerve was isolated from the surrounding connective tissue and separated into the individual branches of the tibialis anterior, extensor digitorum longus, peroneus longus, extensor hallucis longus, and flexor digitorum longus. The nerves innervating each ankle dorsi- and toe flexor muscles were identified via microstimulation of individual nerve branches; all nerve branches except the tibialis anterior were transected.

#### 2.3.2. Plantarflexor Nerve Isolation

The plantarflexor nerve was accessed through the initial incision that was made to access the tibial nerve. The rabbit was positioned with the hind limb in a flexed posture. This position facilitates exposing the tibial nerve by increasing the spacing between muscles of the upper hind limb. Separating the tibial nerve and isolating the individual nerve branches is the most difficult procedure in the surgery. Unlike the peroneal nerve, the tibial nerve is surrounded by blood vessels that must be moved in order to access the nerve. The tibial nerve begins to bifurcate into individual nerve branches proximal to where it dives into the lateral gastrocnemius. The position of initial bifurcation varies greatly between animals with some locations easily accessible and others located beneath muscle and vasculature. Figure 1A demonstrates an ideal scenario where the bifurcation location was easily accessible. When separating the tibial nerve, careful attention must be paid to avoid damaging nerve fibres.

**Figure 1:**
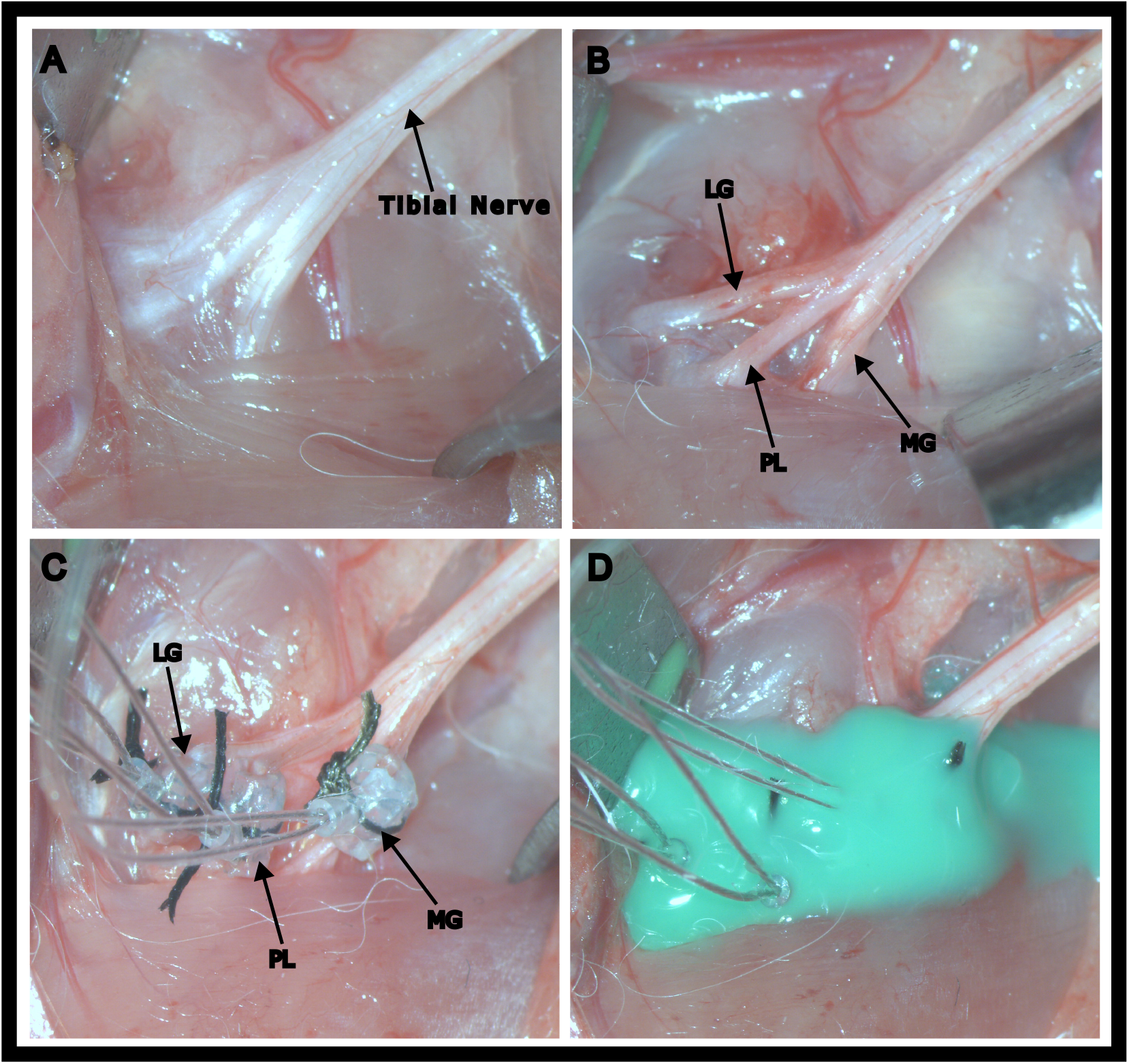
Surgical procedure for the tibial nerve: The tibial nerve is isolated from the surrounding tissue (A). The tibia nerve is separated into the individual nerve branches innervating the gastrocnemius lateralis and soleus, gastrocnemius medialis, and plantaris muscles (B). Once each of the individual nerve branches are isolated, a nerve cuff electrode was wrapped around the nerve and sutured in place (C). A bio-compatible silicone was injected around the nerve branches and each nerve cuff electrode to electrically each electrode from each other and movement of the nerve cuffs (D).

The nerve branches of the PL, MG, and LG+SOL were isolated and a nerve cuff was attached to each (Figure 1C). After attaching the nerve cuff, each muscle was stimulated to ensure full activation of the target muscle - assessed through manual palpation - without concomitant activation of neighbouring muscles. A bio-compatible silicone (Kwik-Cast; World Precision Instruments, Sarasota FL, USA) was applied around each nerve cuff, following instrumentation of each nerve, to prevent unintentional stimulation of neighbouring nerve branches. The incision site was closed and sutured shut to minimize the disturbance to the *in vivo* environment.

### 2.4. Experimental Setup

After the completion of the surgery, rabbits were placed in a supine position and fixed in a custom-built stereotactic frame. Animals was pinned at the left and right iliac crest, medial and lateral left femoral epicondyle, and medial and lateral distal tibial to prevent movement. The animals were positioned with a knee angle of 120 ° and the ankle angle was initially set to 90 °. The rabbit foot was affixed to a rigid foot plate - using tape - that was attached to a servo motor with a one-dimensional load cell to control the position and torque measurement of the ankle joint.

Each nerve cuff was connected to an individual stimulation channel (Stimulator: Grass S8800, Astro/Med Inc., Longueil, QC, Canada). Stimulation voltage was determined with the ankle positioned at 90 °. Individual nerve branches were stimulated at 100 Hz with increasing voltage until a plateau in muscle torque was observed and a further increase in voltage produced no additional torque. Maximum isometric torque was measured at 90 ° of plantarflexion. Muscles were activated using the pre-determined threshold at 100, 60, and 30 Hz stimulation frequency. Stimulation time was set at 1.5 s.

Vital signs (heart rate, breathing clarity/rate, and body temperature) were monitored throughout the experiment and maintained at consistent levels. A hot water pad was placed underneath the rabbit and a heat lamp was located overhead to adjust body temperature, measured via rectal thermometer. Breathing clarity was assessed using a stethoscope, and Glycopyrolate (0.05 mg/kg) was administered in case of obstructed breathing. 40 mL of saline solution was given subcutaneously every four hours to maintain hydration. The isoflurane mixture was adjusted as necessary to maintain heart rate.

At the end of the experiment, individual muscle activation was verified with manual palpation of each muscle during nerve stimulation.

### 2.5. Measurements

Joint torque was measured for each muscle individually and for simultaneous activation of the plantarflexors (PL, LG+SOL, MG) (Figure 2). A three-minute break was included between each trial to prevent muscular fatigue.

**Figure 2:**
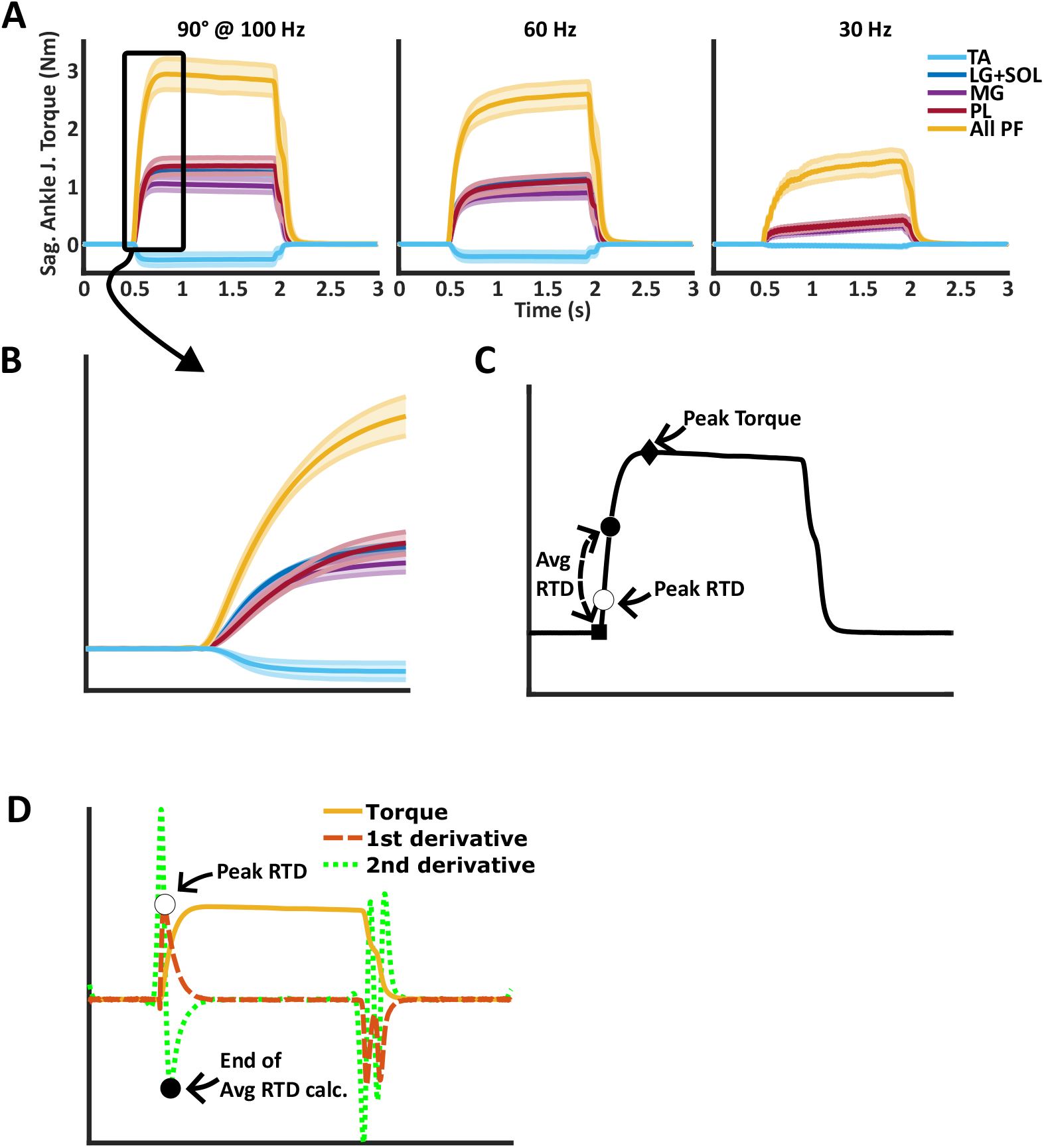
A) Mean and standard error time-series data of sagittal ankle joint torque produced by each individual muscle (LG+SOL = lateral gastrocnemius and soleus, MG = medial gastrocnemius, PL = plantaris, TA = tibialis anterior), and simultaneous activation of all plantarflexor muscles (All PF), for four rabbits at 90° ankle angle and 100, 60, and 30 Hz. B) Highlighting rate of torque development of each muscle at 90° and 100 Hz. C) Peak torque was quantified for each condition (diamond); peak rate of torque development was quantified (white square), and the average rate of torque development was calculated from the onset of contraction (square) to the black circle (Avg RTD). D) Mean torque and the first and second derivative of simultaneous plantarflexor activation. The white circle demonstrates the point at which the peak rate of torque development occurs, and the black circle demonstrates the end point at which the average rate of torque is calculated. The black and white circles in D correspond to the same points in subfigure C.

Peak torque was quantified for all conditions (Figure 2C). Rate of torque development was calculated for each muscle. The peak and average rate of torque development were quantified. Peak rate of torque development was defined as the peak of the first derivative of torque; average rate of torque development was defined as the mean of the first derivative of torque from stimulation onset to the peak of the second derivative of torque - the point at which the decline in the first derivative is greatest (Figure 2D). Linear models were fit to the data, and the effect of stimulation frequency was quantified by computing the t-statistic and associated p-value for the regression coefficients.

To test the reliability of the surgical procedure, each muscle was stimulated three times at 90 ° and 100 Hz, with each stimulation separated by one hour. Linear models were fit to the data, and the effect of time was quantified by computing the t-statistic and associated p-value for the regression coefficient.

Statistical analyses were performed in R (R Core Team, 2018) within RStudio (Version 2021.09.1+372, RStudio, PBC, Boston, MA).

## 3. Results and Discussion

The surgical approach was reliable; a main effect of time was not observed across repeated stimulation of any muscle (*t ≤*1.58, *p≥*0.15).

Mean and standard error ankle joint torque time-series data are presented in Figure 2 for the individual muscle activation and for the simultaneous plantarflexor muscle group activation.

Examples of different muscle properties that can be analysed using this experimental technique are presented in Figure 2. In Figure 2A, mean time-series data for all muscles at each stimulation frequency are presented. The initial rise in torque is highlighted in Figure 2B, demonstrating differences in rate-of-torque development. These differences can be quantified using metrics exhibited in Figure 2C.

When collapsed across all frequencies, peak torque was 0.25 *±* 0.18 Nm for tibialis anterior, 0.97 *±* 0.48 Nm for plantaris, 0.95 *±* 0.38 Nm for lateral gastrocnemius and soleus, 0.76 *±* 0.37 Nm for medial gastrocnemius, and 2.35 *±* 0.82 Nm for simultaneous plantarflexion activation. However, given that torque is known to change with stimulation frequency, linear regression models were fit to the data to determine the effect of frequency on torque time-series metrics. When collapsed across all muscles, peak torque increased by 0.01 Nm for every 1 Hz increase in stimulation frequency (*t* =7.39, *p<*0.001). Peak rate of torque development increased by 0.15 Nm/s and average rate of torque development increased by 0.05 Nm/s for every 1 Hz increase in stimulation frequency (*t ≥*7.00, *p<*0.001).

The experimental setup described herein represents a statically constrained configuration of the lower hind limb in which many muscle properties can be investigated. The utility of this surgical procedure is that it allows muscle activation in isolation or within a muscle group of the lower hind limb. These muscles are critical for locomotion and much of the work on muscle mechanics - particularly, *in vivo* and *in situ* experiments - have been performed on this muscle group; e.g., (Griffiths, 1991; Ryan et al., 2021; Siebert et al., 2015; Tijs et al., 2014; Kelp et al., 2023). Furthermore, given the relatively exposed nature of the Achilles tendon, it may be possible to measure individual muscle forces using tendon buckle force transducers. Understanding how muscles function individually and collectively will help elucidate the importance of position, size, and intermuscular connections of individual muscles to musculoskeletal function.

In conclusion, via the careful isolation of the individual nerve branches of the rabbit lower hind-limb muscles, individual control of the muscles acting at the ankle joint can be achieved. This experimental approach facilitates the investigation of muscle, bone, and joint action in a highly refined scenario that might help expand our understanding of musculoskeletal biomechanics.

## 4. Contributions

The conceptual idea for the project was developed by WH and AS. MB, AS, and SR, developed the surgical technique. MB, AS, and SR performed the data collection. MB performed all data analysis and wrote the initial and final drafts of the manuscript. AS, SR, and WH provided critical feedback on the manuscript. All authors approve of the final version of the manuscript.

